# A Mechanistic Investigation into Ischemia-Driven Distal Recurrence of Glioblastoma

**DOI:** 10.1101/2020.04.03.022202

**Authors:** Lee Curtin, Andrea Hawkins-Daarud, Alyx B. Porter, Kristoffer G. van der Zee, Markus R. Owen, Kristin R. Swanson

## Abstract

Glioblastoma (GBM) is the most aggressive primary brain tumor with a short median survival. Tumor recurrence is a clinical expectation of this disease and usually occurs along the resection cavity wall. However, previous clinical observations have suggested that in cases of ischemia following surgery, tumors are more likely to recur distally. Through the use of a previously established mechanistic model of GBM, the Proliferation Invasion Hypoxia Necrosis Angiogenesis (PIHNA) model, we explore the phenotypic drivers of this observed behavior. We have extended the PIHNA model to include a new nutrient-based vascular efficiency term that encodes the ability of local vasculature to provide nutrients to the simulated tumor. The extended model suggests sensitivity to a hypoxic microenvironment and the inherent migration and proliferation rates of the tumor cells are key factors that drive distal recurrence.

## 1. Introduction

Glioblastoma (GBM) is the most aggressive primary brain tumor [24]. It is uniformly fatal with a median survival from diagnosis of only 15 months with standard of care treatment, consisting of a combination of resection, chemotherapy and radiotherapy [37]. Due to the sensitive location of the tumor, there is a reliance on clinical imaging to assess tumor treatment response and progression. Enhancement on T1-weighted magnetic resonance imaging (T1Gd MRI) with gadolinium contrast shows regions where gadolinium has leaked through disrupted vasculature. T2-weighted MRI (T2 MRI) shows infiltrative edema, fluid that has leaked from vasculature. Abnormalities on T1Gd MRI spatially correlate with the bulk of the tumor mass with central dark regions typically showing necrosis, whereas edema visible on T2 MRI corresponds to regions of lower tumor cell density.

An unfortunate clinical expectation following surgical resection is tumor recurrence, which usually presents on the edge of the resection cavity [9]; this is known as a local recurrence. The recurrent tumor will occasionally enhance on T1Gd MRI in a different region of the brain, away from the primary site, which is known as a distant recurrence [9]. In some cases, the T2 MRI abnormality will become much larger relative to the enhancement on T1Gd MRI, these cases are known diffuse recurrences [9]. In a retrospective study by Thiepold *et al*. it was shown that patients with GBM who had also suffered from perioperative ischemia, defined as an inadequate blood supply to a part of the brain following resection, were more likely to have a distantly and/or diffusely recurring GBM [45]. A disruption in normal vasculature can occur following GBM resection and can lead to ischemia, affecting abnormal tissue in the same way it affects the healthy tissue. By reducing available nutrients to the tumor, the tumor is forced towards a hypoxic phenotype and becomes necrotic if the reduction is sustained. Thiepold attributed the observed difference in recurrence patterns to the hypoxic conditions caused by the reduction in vasculature [45]. In retrospective analyses of patient data, Bette *et al*. found further supporting evidence that perioperative ischemia promoted aggressive GBM recurrence patterns [5,6]. Bette *et al*. showed that perioperative infarct volume was positively associated with more multifocal disease and contact to the ventricle, which have both been shown to negatively impact patient survival in a pretreatment setting [1,35].

Spatiotemporal mathematical models have been used extensively to describe the growth of GBM. These models incorporate features of tumor cells such as cell phenotype, migration, proliferation and interactions with other cells to understand how these influence observed growth behavior in GBM. Such models have the ability to provide mechanistic insight into observed tumor growth patterns and treatment effects. Mathematical models have been created to simulate GBM growth on varying spatial scales from clusters of cells [15,25,36] and murine models [16,22,32] to tissue-level scales seen throughout the presentation of the disease in patients [18,38,39,41,42,44]. Various treatments for GBM have also been modeled, such as resection [27,44], chemotherapy [2,4,7] and radiotherapy [8,22,30], which are all elements of the current standard of care. Other less widely used and experimental treatments have also been modeled such as anti-angiogenic drugs [18,33] and oncolytic virus therapy [13].

An example of a tissue-level growth model of GBM is the Proliferation Invasion Hypoxia Necrosis Angiogenesis (PIHNA) model, which has been used to study different mechanisms of tumor development and shows similar growth and progression patterns to those seen in patient tumors [43]. Simulated hypoxic events have shown an increase in glioma growth rates in spatiotemporal models of GBM [26,28]. We have recently found the parameters of the PIHNA model that drive faster outward growth of these simulated tumors and found that those relating to hypoxia were in some cases extremely influential [12].

To the best of our knowledge, the impact of perioperative ischemia on recurrence in GBM has not been mathematically modeled before. We aim to use the PIHNA model to gain insight into the tumor kinetics that may play a role in this behavior.

In this work, we extend a term in the PIHNA model known as the vascular efficiency term, which determines the ability of local vasculature to provide nutrients to the tumor. We carry this out through the inclusion of a nutrient-transport equation parametrized through glucose uptake rates in GBM. We apply this extended PIHNA model to a set of simulated perioperative ischemia cases to determine influential mechanisms in the model that could drive ischemia-induced distal recurrence patterns in GBM. Specifically, we vary migration and proliferation rates of normoxic and hypoxic cell phenotypes, as well as switching rates between these phenotypes. We find that simulated tumors with faster migration and slower proliferation rates are more likely to recur distantly in cases of perioperative ischemia. We see that this can be promoted by changes in switching rates between normoxic and hypoxic cell phenotypes. We have also simulated these same cases with a less intense ischemic event, and show that this in turn leads to less distantly recurring tumors. Following an initial exploration of simulated resection and ischemic injury, we present a second case example of simulated perioperative ischemia in which we observe similar recurrence patterns that depend on migration and proliferation rates of the normoxic cells. We also present two example simulations, one with and one without ischemia, to show that ischemia can offset the growth of dense tumor in simulations, which may contribute to diffuse recurrence following perioperative ischemia.

## 2. Methods

### 2.1 The PIHNA Model

To simulate glioblastoma growth and spread, we have adapted a previously established tumor growth model – the PIHNA model [12,43]. The PIHNA model describes the growth of GBM with the interactions of vasculature in the process of angiogenesis. This model simulates five different species. that all depend on space and time, and their interactions:

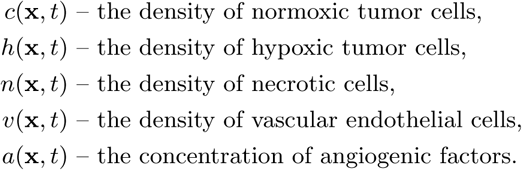

Normoxic cells proliferate with rate *ρ* and migrate with rate *D*_*c*_, whereas hypoxic cells do not proliferate and migrate with rate *D*_*h*_. Cells convert from normoxic to hypoxic phenotypes (with rate *β*) and from hypoxic to normoxic phenotypes (with rate *γ*) depending on the ability of the local vascular density to provide nutrients at their location; hypoxic cells in the model become necrotic if they remain in a vasculature-poor region with rate *α*_*h*_. When any other cell type meets a necrotic cell, they become necrotic with rate *α*_*n*_, as necrotic cells have been shown to encourage cell death through the creation of an unfavorable microenvironment [29,48]. Angiogenic factors migrate with rate *D*_*a*_, are created by the presence of normoxic and hypoxic tumor cells (with rates *δ*_*c*_ and *δ*_*h*_, respectively), decay naturally (*λ*) and are consumed through the creation and presence of vascular cells. For a more in depth justification of these model parameters, see Curtin et al. and Swanson et al. [12,43]. The parameter values used in this work, as well as their units, can all be found in Table 1.

**Table 1:** Parameter definitions and values for the PIHNA model. ^*\**^Parameters that we have added/altered from the original publication of the PIHNA model [43].

|  | Definition | Value/Range | Units | Source |
| --- | --- | --- | --- | --- |
| $D_c$ | Diffusion rate of normoxic cells | $1 - 1000$ | $\frac{\text{mm}^2}{\text{year}}$ | [17] |
| $D_h$ | Diffusion rate of hypoxic cells | $(0.1 - 100)D_c$ | $\frac{\text{mm}^2}{\text{year}}$ | [17, 26]* |
| $\rho$ | Proliferation rate of normoxic cells | $10 - 100$ | $1/\text{year}$ | [17] |
| $\beta$ | Switching rate from normoxia to hypoxia | $0.1\rho, 0.5\rho$ | $1/\text{year}$ | [43]* |
| $\gamma$ | Switching rate from hypoxia to normoxia | $0.005, 0.05, 0.5$ | $1/\text{day}$ | [43] |
| $\alpha_h$ | Switching rate from hypoxia to necrosis | $0.1\beta$ | $1/\text{year}$ | [43]* |
| $\alpha_n$ | Rate of contact necrosis | $\log(2)/50$ | $1/\text{day}$ | [31] |
| $D_v$ | Diffusion rate of endothelial cells | $0.18$ | $\frac{\text{mm}^2}{\text{year}}$ | [43] |
| $D_a$ | Diffusion rate of angiogenic factors | $3.15$ | $\frac{\text{mm}^2}{\text{year}}$ | [43] |
| $\delta_c$ | Normoxic cell production rate of angiogenic factors | $2.77 \times 10^{-13}$ | $\frac{\mu\text{mol}}{\text{cell} \times \text{year}}$ | [43] |
| $\delta_h$ | Hypoxic cell production rate of angiogenic factors | $5.22 \times 10^{-10}$ | $\frac{\mu\text{mol}}{\text{cell} \times \text{year}}$ | [43] |
| $\mu$ | Angiogenesis vasculature production rate | $\log(2)/15$ | $1/\text{day}$ | [43] |
| $q$ | Consumption of angiogenic factors per cell | $1.66$ | $\mu\text{mol}/\text{cell}$ | [43] |
| $\lambda$ | Natural decay rate of angiogenic factors | $15.6$ | $1/\text{day}$ | [43] |
| $\omega$ | Rate of removal of angiogenic factors by vasculature | $\lambda/v_0$ | $\frac{1}{\text{cell} \times \text{day}}$ | [43] |
| $K$ | Maximal cell density | $2.39 \times 10^5$ | $\text{cells}/\text{mm}^3$ | [43] |
| $P_c^w$ | Glucose consumption ratio for normoxic cells in white matter | $1.66 - 4.5$ | - | [14]* |
| $P_h^w$ | Glucose consumption ratio for hypoxic cells in white matter | $1.66 - 4.5$ | - | [14]* |
| $P_c^g$ | Glucose consumption ratio for normoxic cells in grey matter | $0.5 - 2$ | - | [14]* |
| $P_h^g$ | Glucose consumption ratio for hypoxic cells in grey matter | $0.5 - 2$ | - | [14]* |

Following the literature [40], we have assumed that a high total relative cell density of at least 80% is visible on a T1Gd MRI through the aforementioned imaging abnormalities of enhancement and necrosis present on the image. We have also assumed a total relative density of at least 16% is visible on a T2 MRI, due to the spatial correlation between lower cell densities and edema. In the PIHNA model, this translates to the total cell density *T ≥* 0.8 being visible on a T1Gd MRI and *T ≥* 0.16 being visible on a T2 MRI. By construction, the T1Gd lesion is always less than or equal in size to the T2 lesion, which agrees with patient data [17].

We present a schematic for the PIHNA model in Figure 1 that indicates the migration and proliferation of individual species as well as the interactions between all model species. The PIHNA model itself is presented in Equations (1)-(5), which we have annotated to give a full description of each of the terms in the model.

**Fig. 1:**
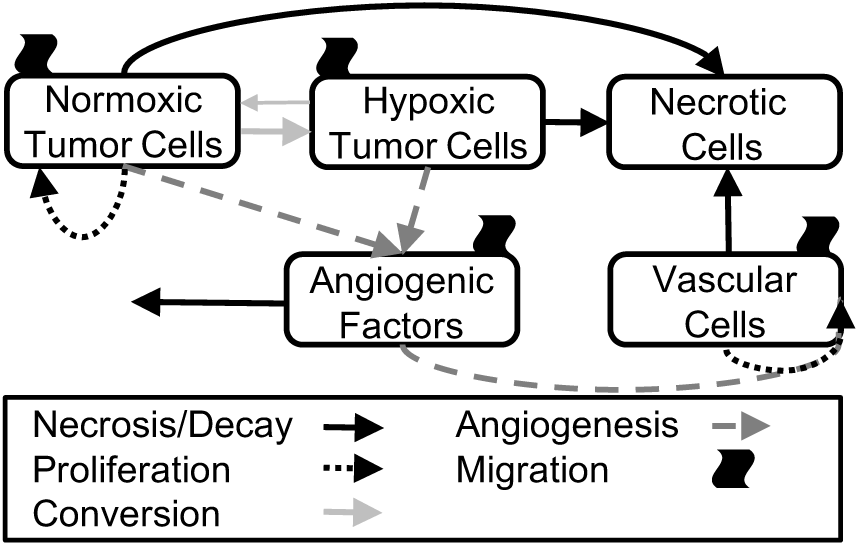
A schematic for the PIHNA model. Normoxic tumor cells (*c*) proliferate, migrate, convert towards hypoxia and can become necrotic. Hypoxic tumor cells (*h*) migrate and can convert back to normoxic cells or to necrotic cells. Necrotic cells (*n*) accumulate as other cell types die. Angiogenic factors (*a*) are created in the presence of normoxic and hypoxic cells, migrate, decay and promote the local creation of vasculature. Vascular cells (*v*) proliferate through the facilitation of angiogenic factors and migrate.

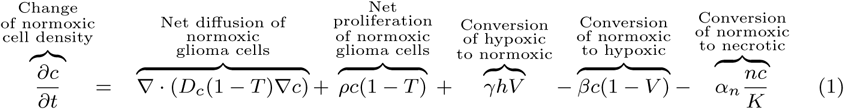

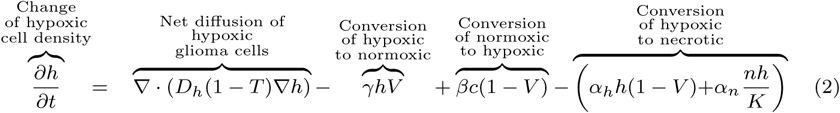

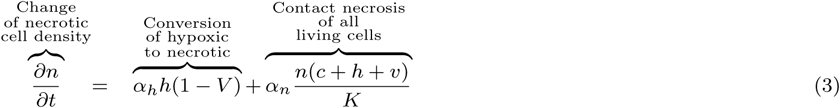

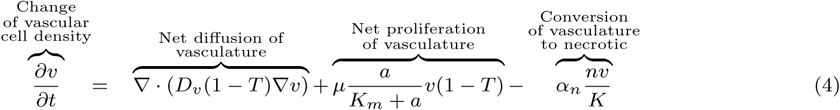

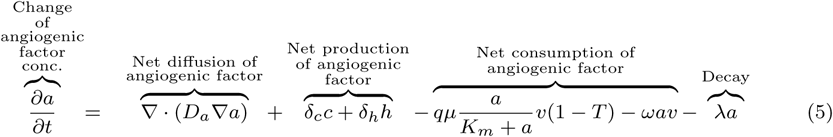

where

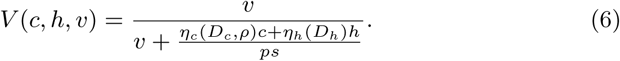

and

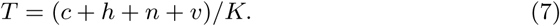

The term *V* is called the vascular efficiency and it models the relationship between the vasculature and its effect on the tumor. We let *V* take values in [0,1] such that it affects the switching rates between the normoxic (*c*), hypoxic (*h*) and necrotic (*n*) cell populations. When vasculature is abundant relative to other cells, *V* is close to 1 representing ample nutrient supply. Whereas when vasculature is relatively low, *V* is close to 0, which represents an unfavorable microenvironment of limited nutrient supply; this promotes conversion towards hypoxic and necrotic cells. In this work, we have extended the vascular efficiency term from previous iterations of the PIHNA model and present the derivation of this term in the next section.

The model equations are run on a two-dimensional slice of a realistic brain geometry from the Brainweb Database [10,11, 20, 21], which spatially differentiates physiological structures such as white matter, grey matter, cerebrospinal fluid (CSF) and anatomical boundaries of the brain. This geometry is an average of multiple MR scans on a single patient to create a brain geometry with 1mm accuracy on and between MR slices. This gives a voxel volume of 1mm^3^, which we use to track tumor volume on the two-dimensional brain slice. The simulations are implemented on white and gray matter, with differences in initial vasculature density and nutrient consumption ratios between these tissues, not allowing for growth of the tumor into the CSF or past the boundaries of the brain.

We initiate the simulation with a small normoxic cell population that decreases spatially from a point with coordinates (*x*_0_, *y*_0_)

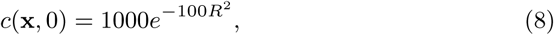

where *R*^2^ = (*x − x*_0_)^2^ + (*y − y*_0_)^2^. The initial seeding locations are described in Section 2.3 and can be seen for the first location in Figure 2.

**Fig. 2:**
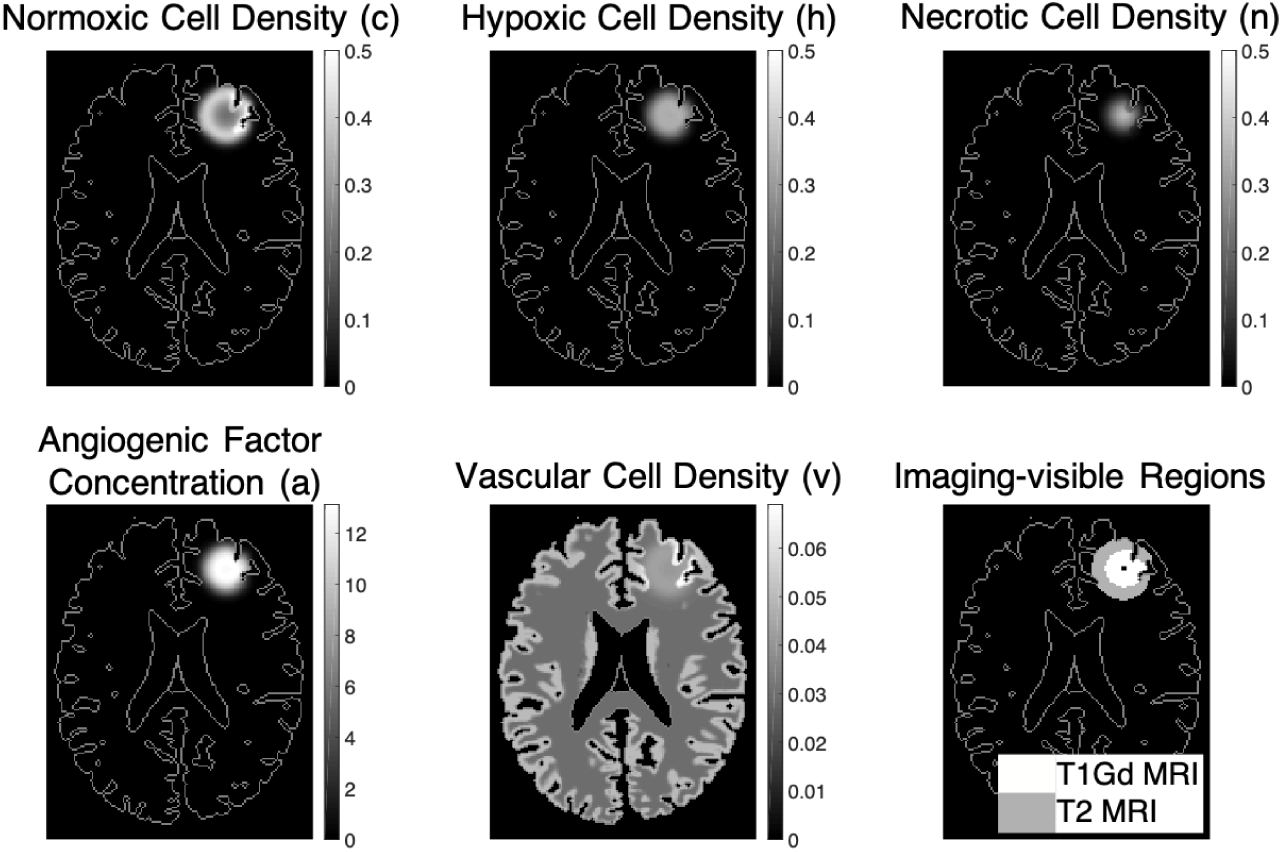
An example simulation shown at a size equivalent to a circle of 1cm radius on simulated T1Gd MRI. We show all cell densities divided by *K* and the angiogenic factor concentration divided by *K*_*M*_. We see how normoxic cells lead the outward growth of the simulated GBM, followed by hypoxic cells and necrotic cells. Angiogenic factors are mostly found in the hypoxic cell region. We also show the regions that are assumed visible on T1Gd MRI (*T ≥* 0.8) and T2 MRI (*T ≥* 0.16) as well as the point where the tumor is initiated (black pixel). In this simulation, *D*_*h*_*/D*_*c*_ = 10, *D*_*c*_ = 10^1.5^mm^2^/year, *ρ* = 100/year, *β* = 0.5*ρ* and *γ* = 0.05/day.

The initial vascular cell densities are heterogeneous, set to 3% and 5% of the carrying capacity, *K*, in white and grey matter respectively; these values fall within the values for cerebral blood volume found from the literature [47]. All other spatiotemporal variables are initially set to zero. There are no-flux boundary conditions on the outer boundary of the brain for all variables as well as on the CSF that do not allow growth outside of the brain or into CSF regions.

### 2.2 Nutrient-Based Vascular Efficiency

The extended vascular efficiency term uses a reaction-transport equation to model the nutrient consumption by the tumor cells. Using this reaction-transport equation for the movement and consumption of nutrient, *f* ^1^, the derivation of the vascular efficiency term goes as follows:

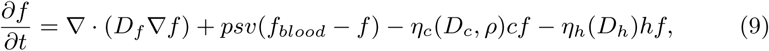

where *p* is the permeability of the blood brain barrier to nutrient, *s* is the vascular surface area per unit volume, *f*_*blood*_ is the concentration of nutrient in the blood which is assumed fixed^2^, *η*_*c*_ is the rate of nutrient consumption by normoxic cells and *η*_*h*_ is the nutrient consumption rate by hypoxic cells. We have let *η*_*c*_ depend on the diffusion (*D*_*c*_) and proliferation (*ρ*) rates of normoxic cells, as these processes require energy. A larger *D*_*c*_ and *ρ* will require more energy as the tumor cells migrate and proliferate relatively quickly. Similarly, we have set *η*_*h*_ to depend on the value of *D*_*h*_, as faster migrating tumor cells require more energy and in turn more nutrient.

Now if we assume that in the timescale of interest, the nutrient concentration rapidly reaches steady state, and that the nutrient is consumed much faster than it diffuses, we can eliminate those terms to be left with

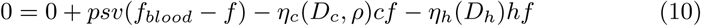

and rearrange to get

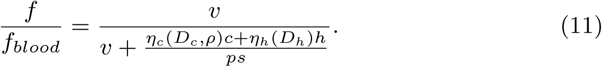

We assign this expression as the vascular efficiency term, *V*, as it corresponds to the ability of the vasculature to provide nutrients to the tumor. This term is similar to that seen in the original formulation of the PIHNA model but now includes the nutrient consumption and extravasation of nutrients from the blood [43].

To estimate the parameters *η*_*c*_, *η*_*h*_ and *ps*, we used Fludeoxyglucose (FDG) Positron Emission Tomography (PET) data from a paper by Delbeke *et al*. [14]. FDG is analogous to glucose and can be detected on PET scans. We have chosen glucose as an estimate for our generic nutrient due to the availability of imaging data that we could use to parametrise our vascular efficiency term. Due to the increase in anaerobic respiration of cancer cells compared with normal tissue, known as the Warburg effect [23,46], we might expect oxygen uptake ratios to be lower than glucose.

We note that to parametrize our nutrient-based vascular efficiency term, we only need to consider the ratio between *η*_*c*_:*ps* and *η*_*h*_:*ps*. As both of these expressions are in the same units of mm^3^*/*cell*/*year, their ratio is dimensionless. Delbeke presents the uptake ratios between tumor and healthy tissue within both white and grey matter. To make use of these values, we assume that in a homeostatic healthy brain, the rate of glucose being used by healthy tissue that is not vasculature is equal to the rate of glucose entering from the vasculature. We do not, however, model healthy tissue in the current formulation of the PIHNA model. For the benefit of this section, let us introduce unaffected healthy tissue *u*_0_, with glucose uptake rate *η*_*u*_, we assume

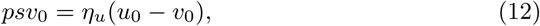

where *v*_0_ is the initial background vascular cell density in the PIHNA model, and *u*_0_ is the healthy tissue density. We then have *ps* = *η*_*u*_(*u*_0_*/v*_0_ *−*1), which will always be positive in PIHNA simulations as vasculature takes up a small percentage of brain volume compared to other tissue. We assume that in healthy white matter tissue there is 3% vasculature and in grey there is 5%, so we let *v*_0_*/u*_0_ = 0.03 in white matter and *v*_0_*/u*_0_ = 0.05 in grey matter; these values fall within realistic values for cerebral blood volume [47]. Now the ratios of glucose uptake rates by tumor to the glucose uptake rates by healthy tissue given by Delbeke can be considered as various values of *P*_*c*_ = *η*_*c*_*/η*_*u*_ and *P*_*h*_ = *η*_*h*_*/η*_*u*_ in the PIHNA model. So Equation 11 is now expressed as

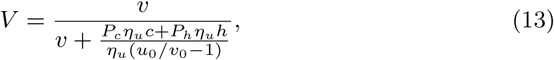

and the *η*_*u*_ terms cancel to give

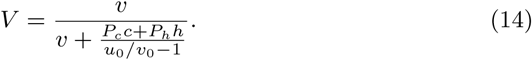

We noted that in the work by Delbeke et al.[14] there was a spread of relative tumor uptake values for high grade gliomas within cortical and white matter tissue across 20 patients. As an approximation, we attributed these differences to the nutritional demands of the individual high grade gliomas. We assign normoxic cells with high (low) *D*_*c*_ and high (low) *ρ* in the PIHNA model with the higher (lower) glucose uptake rates from the literature, which also vary between white and grey matter. We assign hypoxic cells with high (low) *D*_*h*_ high (low) glucose uptake rates in the same manner as the normoxic cells. The values in between the extremes are assigned using a log linear scale, due to the large range of *D*_*c*_, *D*_*h*_ and *ρ* values used in PIHNA simulations. The ratios 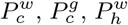 and 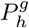 are then given by

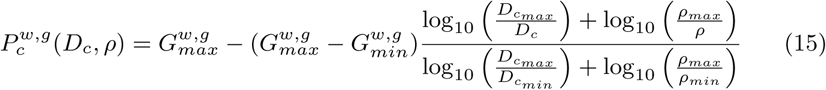

and

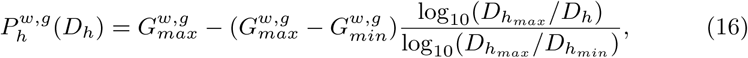

where we use the extremes of tumor to normal tissue uptake ratios in white matter (taken as 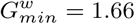 and 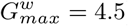) and the extremes of observed uptake ratio in grey matter (taken as 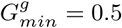 and 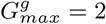) as the minimum to maximum glucose uptake ratios *G*_*min*_ and *G*_*max*_. The maximum and minimum *D*_*c*_, *D*_*h*_ and *ρ* values are equal to the maximal and minimal rates that we run in our simulations, see Table 1. This along with the values of *v*_0_*/u*_0_ give the parametrisation of the nutrient-based vascular efficiency term.

### 2.3 Modeling Resection and Ischemia

Using the PIHNA model, we have simulated a resection that occurs once the tumor has grown to a shape with a volume equivalent to a disc of 1cm radius on simulated T1Gd imaging. The tumor was initiated at *x*_0_ = 100, *y*_0_ = 150 on the 85th axial slice of the brain geometry. Post resection, zero-flux boundary conditions are added around the simulated frontal lobectomy (green outline in Figure 3) so that regrowth into the resection cavity is not possible. This boundary condition was chosen as recurrence into the resection cavity itself is unlikely in GBM. GBM cells have a migratory nature and need structure and nutrients to migrate and grow [3]. As there is a lack of tissue and vasculature within the resection cavity, it is not an environment conducive to GBM growth. Every resection is the same, in that the same region of brain geometry is removed, which removes all of the enhancing T1Gd region. To incorporate the potential reality that surgery could induce a nearby ischemic event (red outline in Figure 3), we add subsequent ischemia through a transient reduction in the vasculature term, *v*, to a region adjacent to the resection cavity wall. We have modeled ischemia as a reduction only at the time point of resection, the vessels then continue to follow the model equations. We reduced the vasculature to 1% of its value at the time of resection thus simulating a near complete ischemic event in the region noted in red in Figure 3, which is also the value used in all recurrence pattern figures presented in the main text. In further simulations, we have also reduced the vasculature to 10% to see the impact of this less intense simulated ischemia on recurrence locations; results of these simulations are presented in Appendix Figures 12 - 14.

**Fig. 3:**
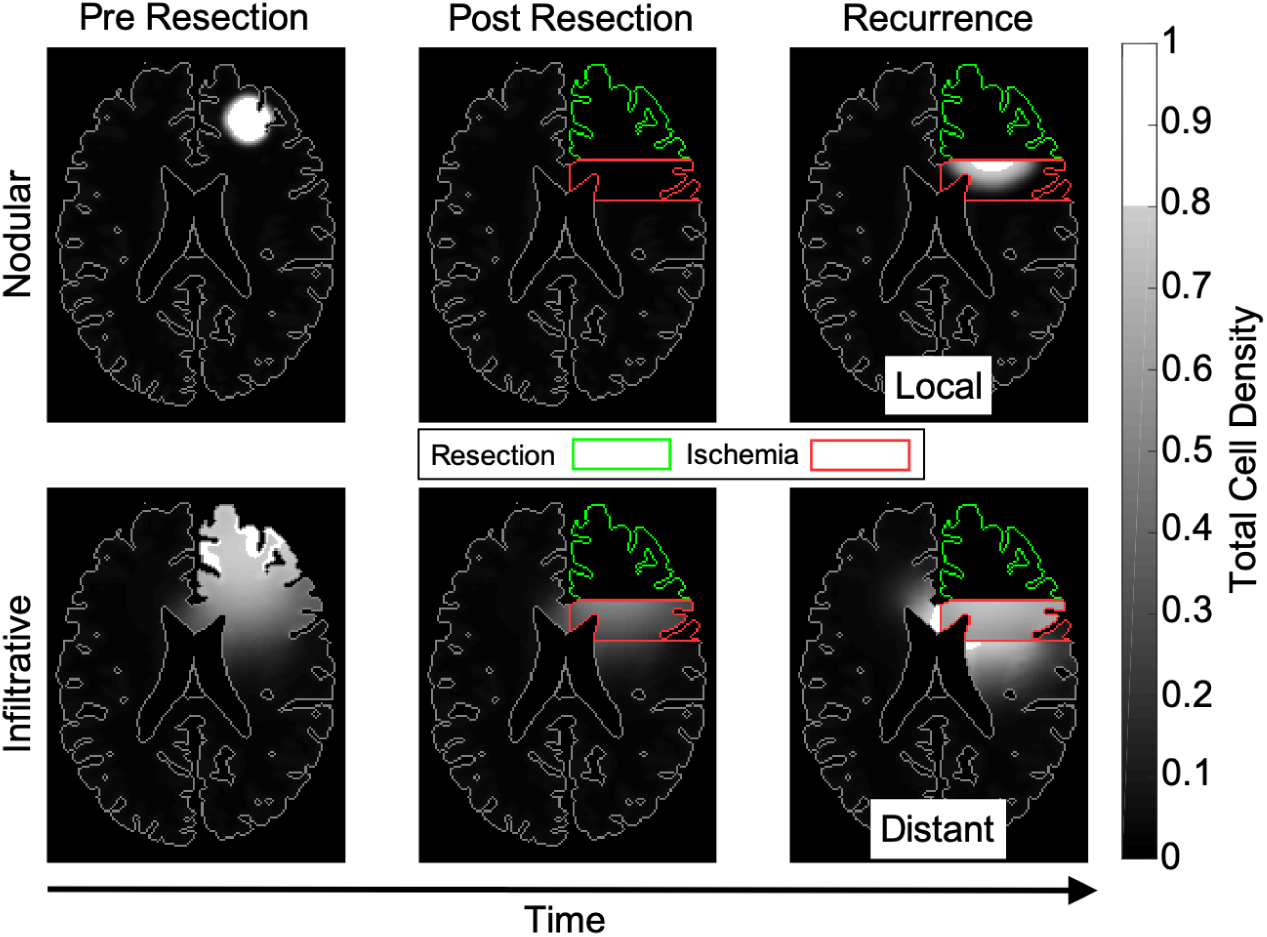
The tumor undergoes resection that removes the T1Gd imageable tumor cell density at 1cm radius (assumed at 80% of the maximum cell density and shown in white) as well as the surrounding tissue. In these two examples, the nodular tumor (top row) recurs locally, whereas the infiltrative tumor (bottom row) recurs distantly. In these simulations, *β* = 0.5*ρ, γ* = 0.05/day and *D*_*h*_ = 10*D*_*c*_. For the nodular tumor, *D*_*c*_ = 10^0.5^mm^2^/year and *ρ* = 10^1.5^/year. For the infiltrative tumor, *D*_*c*_ = 100mm^2^/year and *ρ* = 10/year.

We also present a second location with a recurrence that occurs when the tumor has reached a volume of a disc of 0.25cm radius on T1Gd imaging. In this setting, the tumor location was initiated at *x*_0_ = 44, *y*_0_ = 132 on the 89th coronal slice of the brain geometry. This resection cavity was simulated as a 10 pixel radius around the initial tumor location, with ischemia as a further 5 pixel radius. Zero flux boundary conditions are added to this resection cavity and perioperative ischemia is simulated in the surrounding tissue.

### 2.4 Virtual Experiments

We run simulations for different values of normoxic cell migration (*D*_*c*_ with range 1 *−* 1000mm^2^ /year) and proliferation (*ρ* with range 10 *−* 100/year), as well as test two values of *β* (0.1*ρ* and 0.5*ρ*), which is the switching rate from the normoxic cell density towards hypoxic cell density, three values of *γ* (0.005, 0.05, 0.5/day), which is the switching rate back from hypoxic cell density to a normoxic cell density, and the rate of hypoxic to normoxic cell migration, *D*_*h*_*/D*_*c*_ (1, 10 or 100). We vary the ratio of hypoxic to normoxic cell migration due to evidence that GBM cells migrate faster in hypoxic conditions [19,49]. These parameters were chosen as they represent the tumor’s response to hypoxic stress. In previous work, we have observed that all of these parameters (except for *β*) influence the outward growth rate of PIHNA simulations, which is another consideration of the effects of hypoxia on GBM [12]. We note that the varying tumor kinetics (migration rates *D*_*c*_, *D*_*h*_ and proliferation rate *ρ*) affect the nodularity of the simulated tumors. Simulations with higher ratios of migration to proliferation will be more infiltrative tumors, whereas those with higher ratios of proliferation to migration will be denser tumor masses with less infiltration and more well-defined tumor cell density boundaries. Examples of this effect can be seen in Figure 3. We ran further simulations of a second tumor location with *β* = 0.5*ρ, γ* = 0.5/day and *D*_*c*_ = 10*D*_*h*_, as a validation that the observed dependency of recurrence patterns on migration and proliferation rates were not simply a function of the resection and ischemic regions that we chose in the first location.

### 2.5 Defining Recurrence Location

Recurrence location of a tumor is classified as the reappearance of the tumor on T1Gd MR scans as is done clinically [50]. If a tumor initially reappears outside of the simulated ischemic region above a certain thresholded size (a disc of radius 2mm on simulated T1Gd MRI) before appearing anywhere else, it is classified as distant. Whereas if it appears within the ischemic region along the cavity wall above the same threshold before anywhere else, it is classed as a local recurrence. Examples of these cases can be seen in Figure 3. We do not consider the nature of T2 signal in our definition of distant recurrence. We define a mixed recurrence when the tumor appears on simulated T1Gd MRI both inside and outside the ischemic region before the size threshold within either region is reached. This method of defining distant, local and mixed recurrence patterns is applied to both tumor locations presented in this work.

## 3. Results

### 3.1 Individual tumor kinetics affect recurrence location following perioperative ischemia

Extending on the paper by Thiepold *et al*., that suggests distant recurrence can occur through ischemia and subsequent hypoxia [45], the PIHNA model suggests that tumor kinetics also play a role. Figure 3 shows two simulated tumors, one nodular and the other infiltrative, that go through resection and subsequently recur. The recurrence pattern for the nodular tumor is local, whereas the infiltrative tumor recurs distantly. Such distantly recurring tumors remain in lower cell densities within the ischemic region and appear outside on simulated T1Gd imaging as they continue to increase their cell density outside of the ischemic region. The only differences between the two simulations presented in Figure 3 are the migration and proliferation rates of the tumor cells.

### 3.2 Tumor response to hypoxic conditions affects recurrence location

By varying individual tumor kinetics (*D*_*c*_ and *ρ* with *D*_*h*_ = 10*D*_*c*_) and the maximal rate at which tumor cells become hypoxic and in turn necrotic (*β*), with a fixed vascular ischemia post resection, we are able to show differing tumor recurrence locations, see Figure 4. We also varied the maximal rate at which hypoxic tumor cells returned to a normoxic state, *γ*. Changing this parameter also had an effect on the recurrence patterns, see Figure 5. A low level of *γ* promotes more distant recurrence, while a high level of *γ* promotes local recurrence. Recurrence location is classified as the first reappearance of a tumor on T1Gd MR imaging as described in the previous section.

**Fig. 4:**
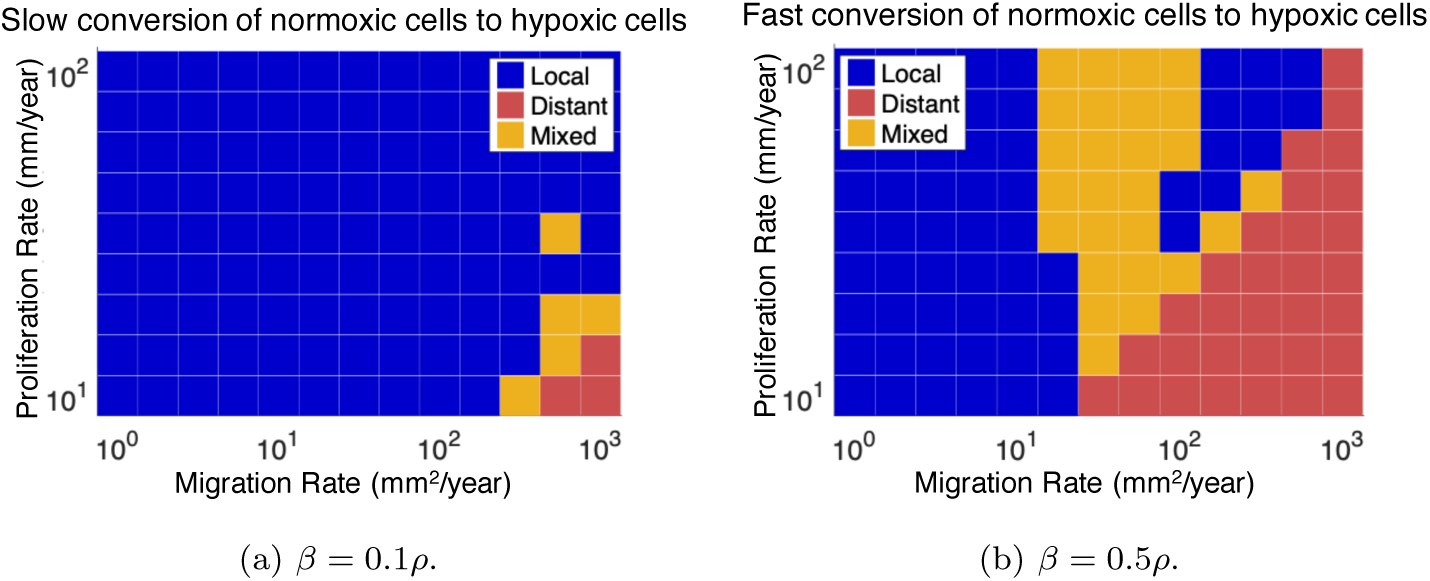
Recurrence location classified for various *D*_*c*_, *ρ* and *β* for *D*_*h*_ = 10*D*_*c*_ and *γ* = 0.05/day. We see that higher values of *β* (the conversion rate from normoxic to hypoxic cells) lead to a larger proportion of distant recurrences in *D*_*c*_ and *ρ* parameter space. Higher migration rates, *D*_*c*_, and lower proliferation rates, *ρ*, lead to more distantly recurring simulated tumors.

**Fig. 5:**
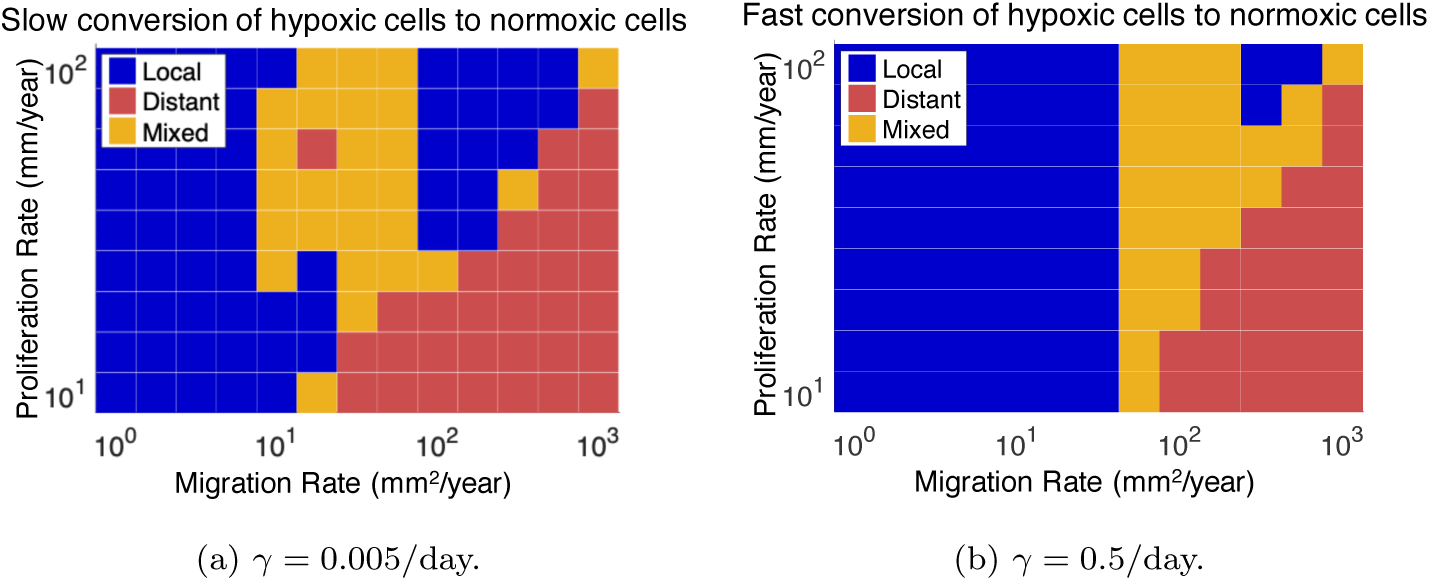
Recurrence location classified for various *D*_*c*_, *ρ* and *γ* for *D*_*h*_ = 10*D*_*c*_ and *β* = 0.5*ρ*. We see the higher (lower) values of *γ* lead to a lower (higher) proportion of distant recurrences in *D*_*c*_ and *ρ* parameter space. Higher migration rates, *D*_*c*_, and lower proliferation rates, *ρ*, lead to more distantly recurring simulated tumors.

An increase in *β* leads to more sensitivity in the tumors to ischemia, which causes them to become more hypoxic and therefore less proliferative within the ischemic region. They are more likely to become more dense, and therefore imageable on simulated T1Gd MRI, outside of the ischemic region and be seen as a distant recurrence. Conversely, an increase in *γ*, the conversion rate from a hypoxic cell phenotype back to normoxic, hinders this effect as it limits the impact of hypoxia on the growth of the simulated tumor. We present other simulation results of varying *β* and *γ* in Appendix Figure 10.

### 3.3 Faster Hypoxic Cell Migration Rates Promote Distant Recurrence

Following this initial analysis, we also varied the hypoxic diffusion rate relative to the normoxic counterpart, *D*_*h*_*/D*_*c*_. Along with the simulations where *D*_*h*_ = 10*D*_*c*_ described in the previous section, we have set *D*_*h*_*/D*_*c*_ = 1 and *D*_*h*_*/D*_*c*_ = 100 (see Figure 6). We see that the higher the *D*_*h*_*/D*_*c*_ value, the more distantly recurring tumors occur for fixed values of *β* and *γ*. The effect of an increase in *D*_*h*_*/D*_*c*_ is more pronounced for tumors that are more sensitive to the hypoxic environment caused by the ischemia, see Appendix Figures 9 - 11. In previous work, we have shown that an increase in *D*_*h*_*/D*_*c*_ increases the outward growth rate of PIHNA-simulated GBM [12]. With faster hypoxic migration rates, the simulated tumor cell densities are able to travel through the hypoxic region faster. These tumors can then reach the region of the brain slice unaffected by ischemia and develop into a dense tumor mass before the tumor develops within the ischemic region.

**Fig. 6:**
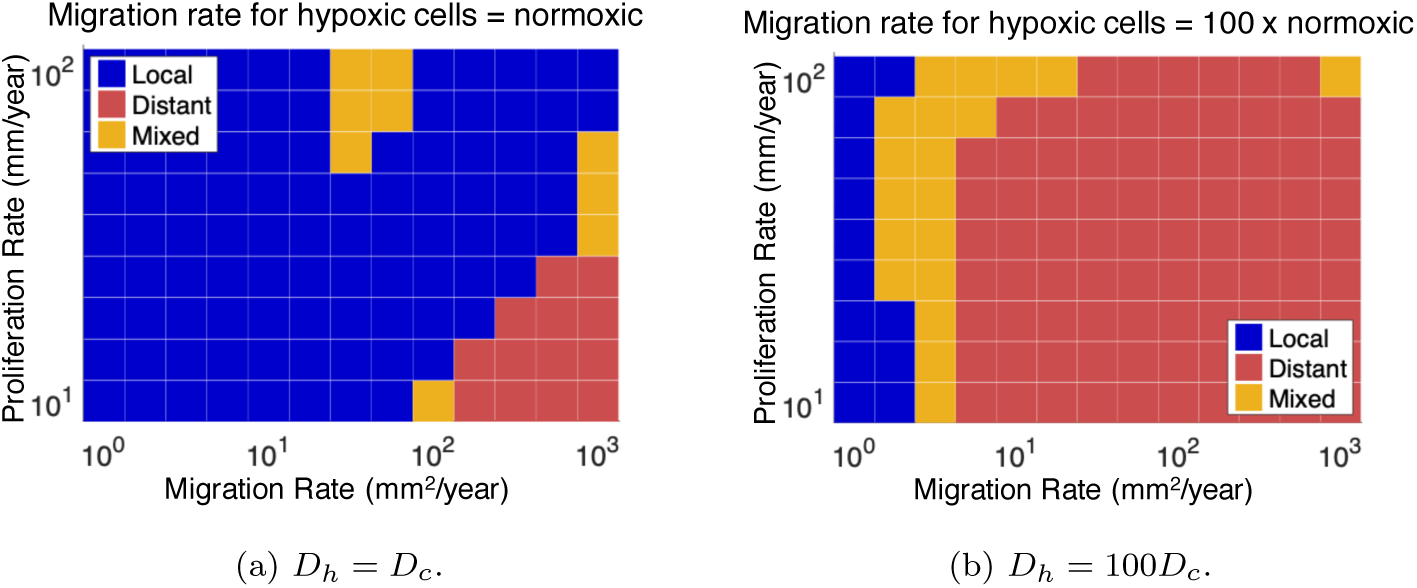
Recurrence location classified for various *D*_*c*_, *ρ* and *D*_*h*_*/D*_*c*_ for *β* = 0.5*ρ*. We see that the higher (lower) level *D*_*h*_*/D*_*c*_ lead to a larger (smaller) proportion of distant recurrences in *D*_*c*_ and *ρ* parameter space. Higher migration rates, *D*_*c*_, and lower proliferation rates, *ρ*, lead to more distantly recurring simulated tumors.

**Fig. 7:**
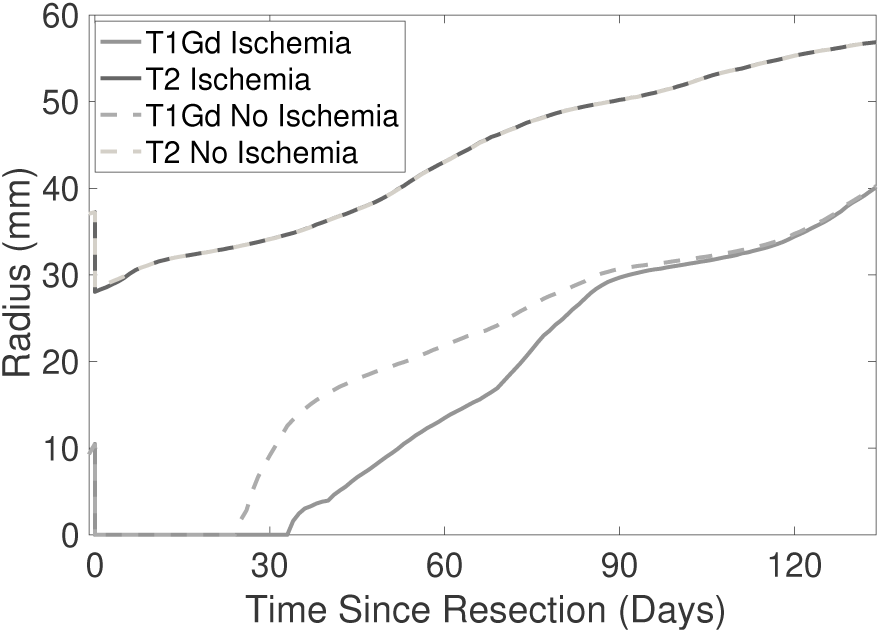
We present the imageable tumor radii as a function of time for two simulations. The only difference between these simulations is that one has perioperative ischemia and the other does not. Note that the growth of the T1Gd imageable radius is delayed by the ischemia, whereas the T2 imageable radius is minimally affected. This offset of dense tumor growth may be a contributing factor to diffuse recurrence following perioperative ischemia. In these simulations, *D*_*c*_ = 10^2.5^mm^2^/year, *ρ* = 10^1.25^/year, *D*_*h*_ = 10*D*_*c*_, *β* = 0.5*ρ* and *γ* = 0.005/day.

**Fig. 8:**
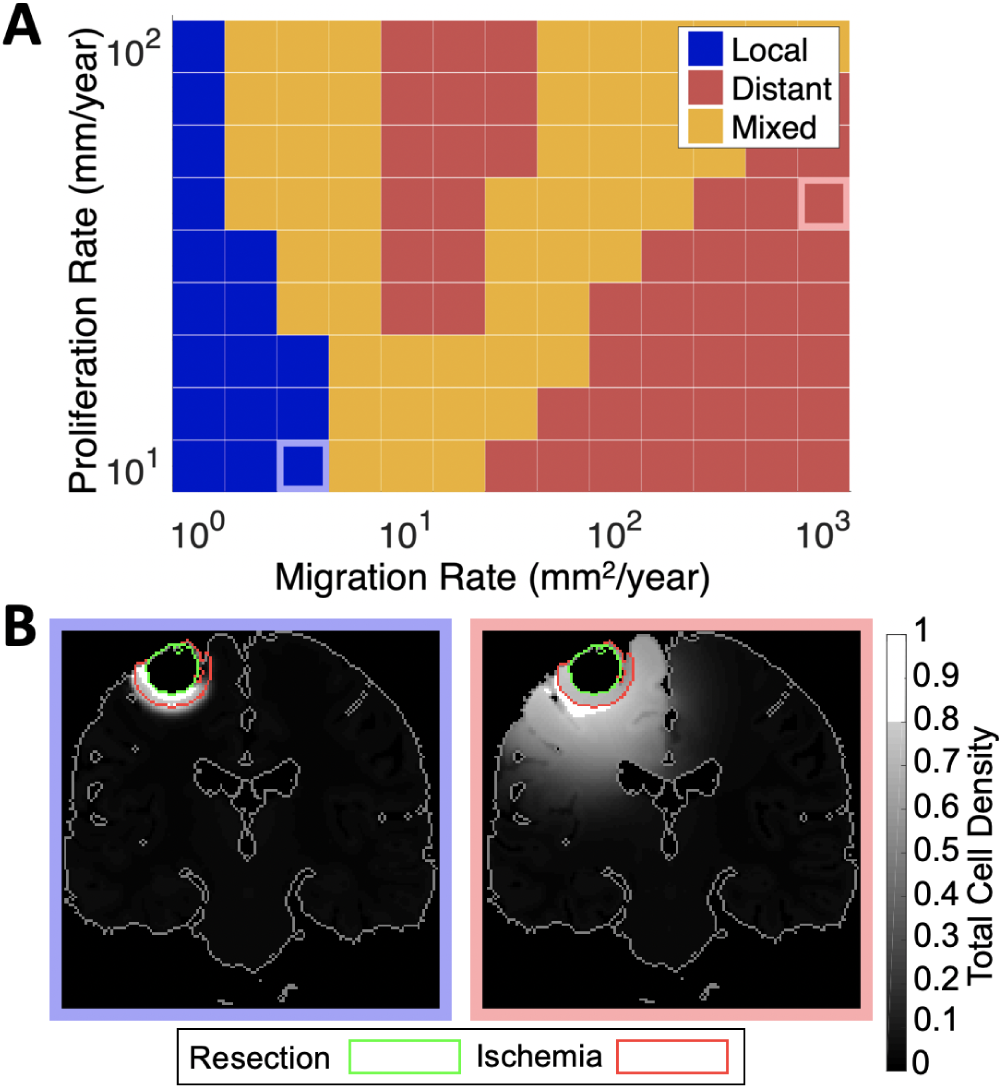
A) Recurrence location classified for the second tumor location in coronal view, for various *D*_*c*_ and *ρ* values. As in the first location, tumors with more diffuse characteristics recur distantly, while those with lower migration rates and faster proliferation rates tended to be more mixed. In these simulations, *β* = 0.5*ρ, γ* = 0.5/day and *D*_*h*_ = 10*D*_*c*_, parameter values are equivalent to Subfigure 5b. B) Two example simulations, with local recurrence (left) and distant recurrence (right), showing the resection (inner green outline) and ischemic (outer red out-line) regions. Migration and proliferation rates of these example simulations are indicated on Subfigure A.

**Fig. 9:**
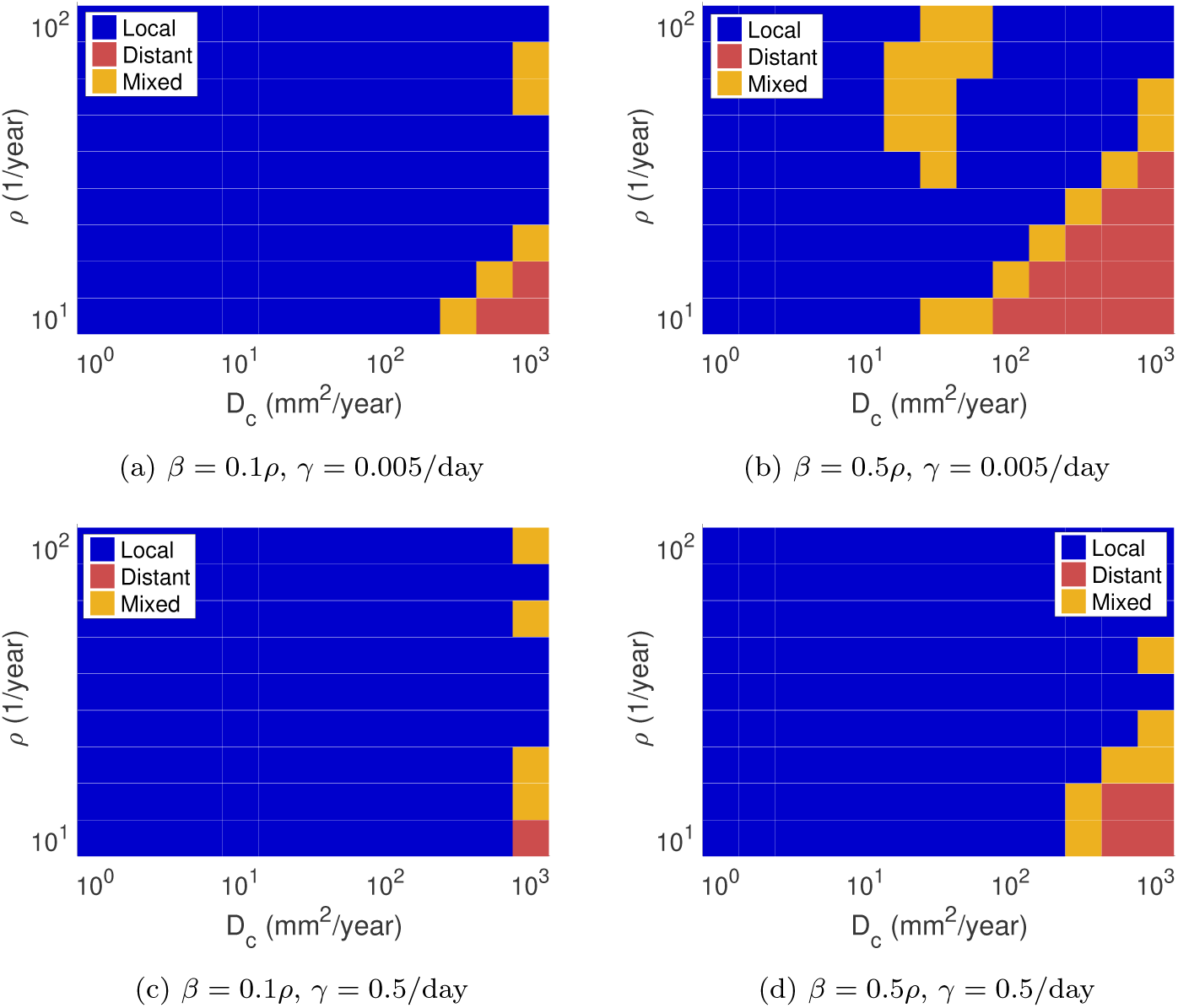
Recurrence location classified for various *D*_*c*_, *ρ, β* and levels of ischemia for *D*_*h*_ = *D*_*c*_ for *γ* = 0.005/day and *γ* = 0.5/day. We see that higher values of *β* and lower levels of *γ* lead to a larger proportion of distant recurrences in *D*_*c*_ and *ρ* parameter space. Higher migration rates, *D*_*c*_, and lower proliferation rates, *ρ*, lead to more distantly recurring simulated tumors.

**Fig. 10:**
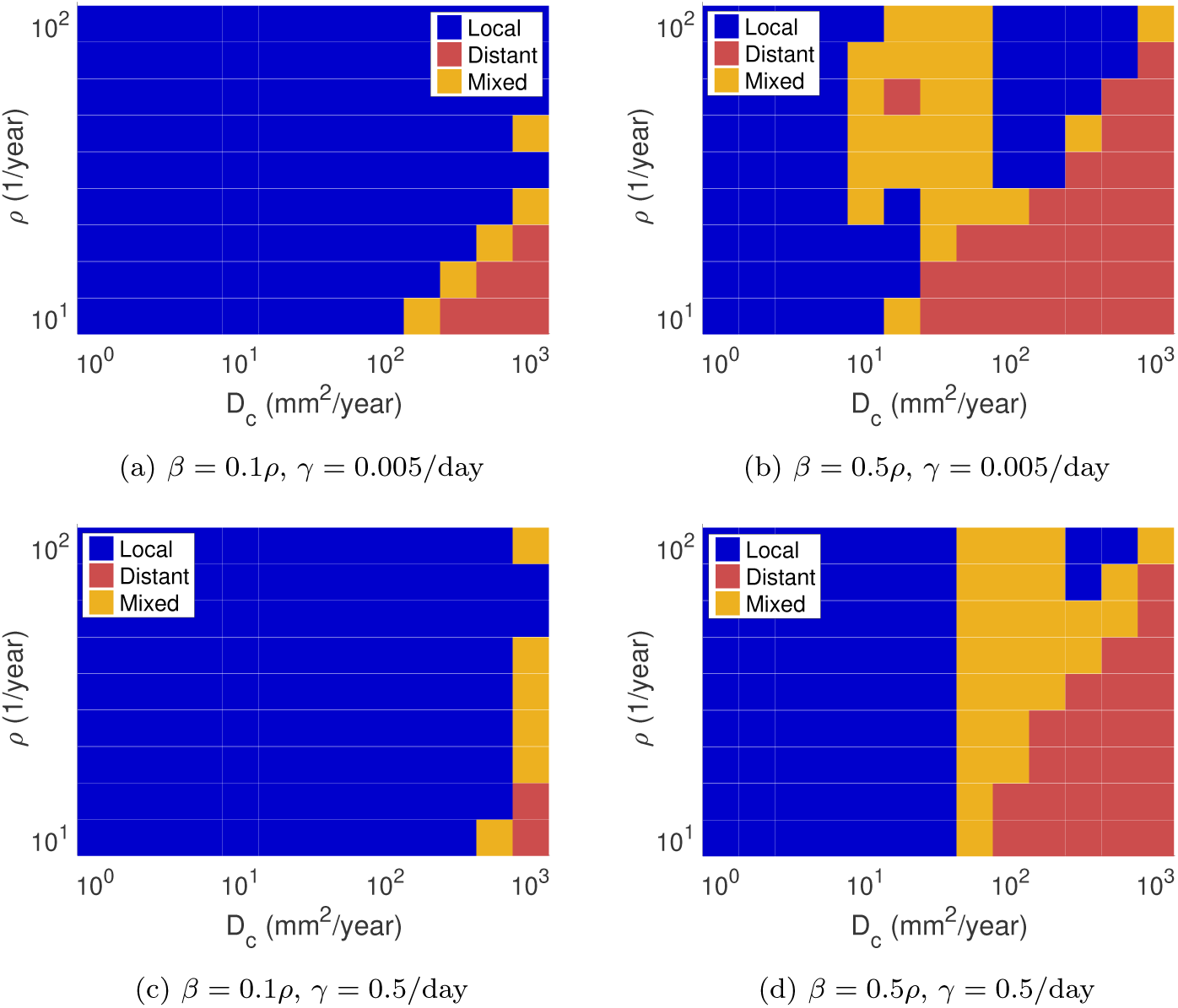
Recurrence location classified for various *D*_*c*_, *ρ, β* and levels of ischemia for *D*_*h*_ = 10*D*_*c*_ for *γ* = 0.005/day and *γ* = 0.5/day. We see that higher values of *β* and lower levels of *γ* lead to a larger proportion of distant recurrences in *D*_*c*_ and *ρ* parameter space. Higher migration rates, *D*_*c*_, and lower proliferation rates, *ρ*, lead to more distantly recurring simulated tumors.

### 3.4 Diffuse Recurrence Present Through an Ischemia-Induced Reduction in Dense Tumor

The paper by Thiepold considered distal recurrence as either diffuse or distant [45]. Diffuse recurrence presents as a marked increase in T2 signal without an accompanying increase in T1Gd enhancement. To explore how diffuse recurrence may occur in the PIHNA model, we present a plot of T1Gd and T2 volume over time for a simulated case of perioperative ischemia, compared with the same simulation without such ischemia. This case recurred distantly and is an example of parameters used in Subfigure 5a, with *D*_*c*_ = 10^2.5^mm^2^/year, *ρ* = 10^1.25^/year, *D*_*h*_ = 10*D*_*c*_, *β* = 0.5*ρ* and *γ* = 0.005/day. As can be seen in Figure 7, we see a reduction in T1Gd signal as a result of the perioperative ischemia for a period of growth, while T2 signal is less affected. If the tumor with perioperative ischemia was observed in this window, the relative T2 signal compared with T1Gd enhancement would appear larger, resulting in a higher probability of the tumor being considered diffuse at recurrence. After some further growth, the T1Gd radii of both simulations meet, as the tumor within the ischemic region eventually grows to a high-enough density to be visible on T1Gd MRI.

### 3.5 Extent of Vasculature Reduction Impacts Recurrence Patterns

PIHNA model simulations were run with a lower level of reduction in functional vasculature to 10% within the same geometry of perioperative ischemia shown in Figure 3. This was implemented for all changes in *β, γ* and *D*_*h*_ presented in the main text. In this setting, all recurrence location results in Figures 4 - 6 showed a shift towards more local recurrence within the values of *D*_*c*_ and *ρ* that were tested. We present these corresponding recurrence location figures in the Appendix (Figures 12 - 14).

### 3.6 Validation in a Second Tumor Location

To explore the simulated recurrence location further, we ran PIHNA simulations for a second tumor location in a coronal view. This tumor was initiated closer to the surface of the brain and resected at a smaller T1Gd imageable tumor volume (equivalent to a disc of 0.25cm radius). We present recurrence locations and example simulations for *β* = 0.5*ρ, γ* = 0.5/day and *D*_*h*_ = 10*D*_*c*_ with varying migration and proliferation rates in Figure 8. Other than the different location and size at resection, these simulations have the same parameters as Subfigure 5b. As we observed in the first location, tumors with more diffuse characteristics recurred distantly, whereas those with more nodular characteristics and lower migration rates tended to recur locally or mixed. In this second location, we observe some simulated tumors with faster proliferation rates but mid-range migration rates that also recur distantly, which we also saw some evidence of in the first location, present in Subfigures 4b, 5a and 6a.

**Fig. 11:**
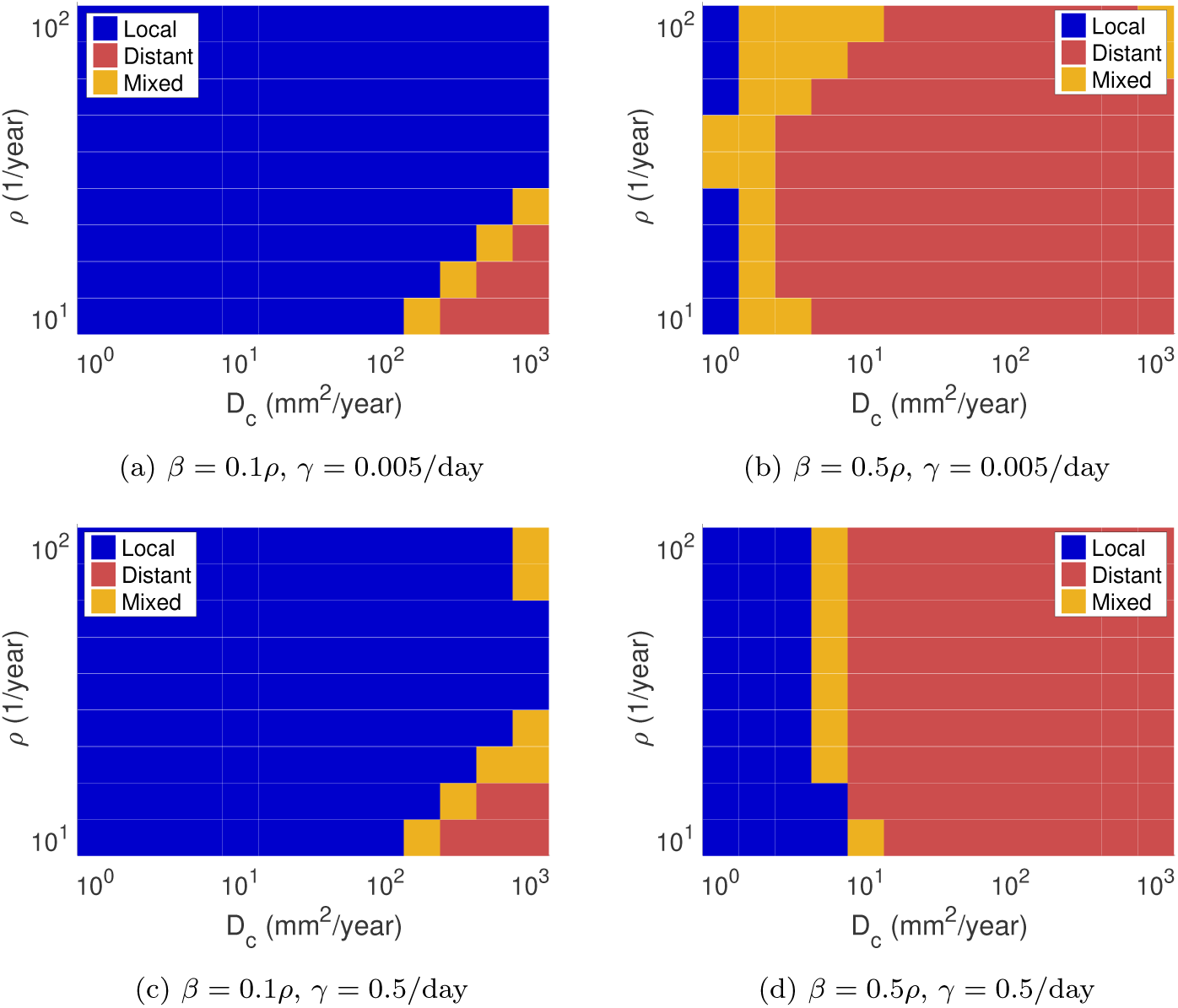
Recurrence location classified for various *D*_*c*_, *ρ, β* and levels of ischemia for *D*_*h*_ = 100*D*_*c*_ for *γ* = 0.005/day and *γ* = 0.5/day. We see that higher values of *β* and lower levels of *γ* lead to a larger proportion of distant recurrences in *D*_*c*_ and *ρ* parameter space. Higher migration rates, *D*_*c*_, and lower proliferation rates, *ρ*, lead to more distantly recurring simulated tumors.

**Fig. 12:**
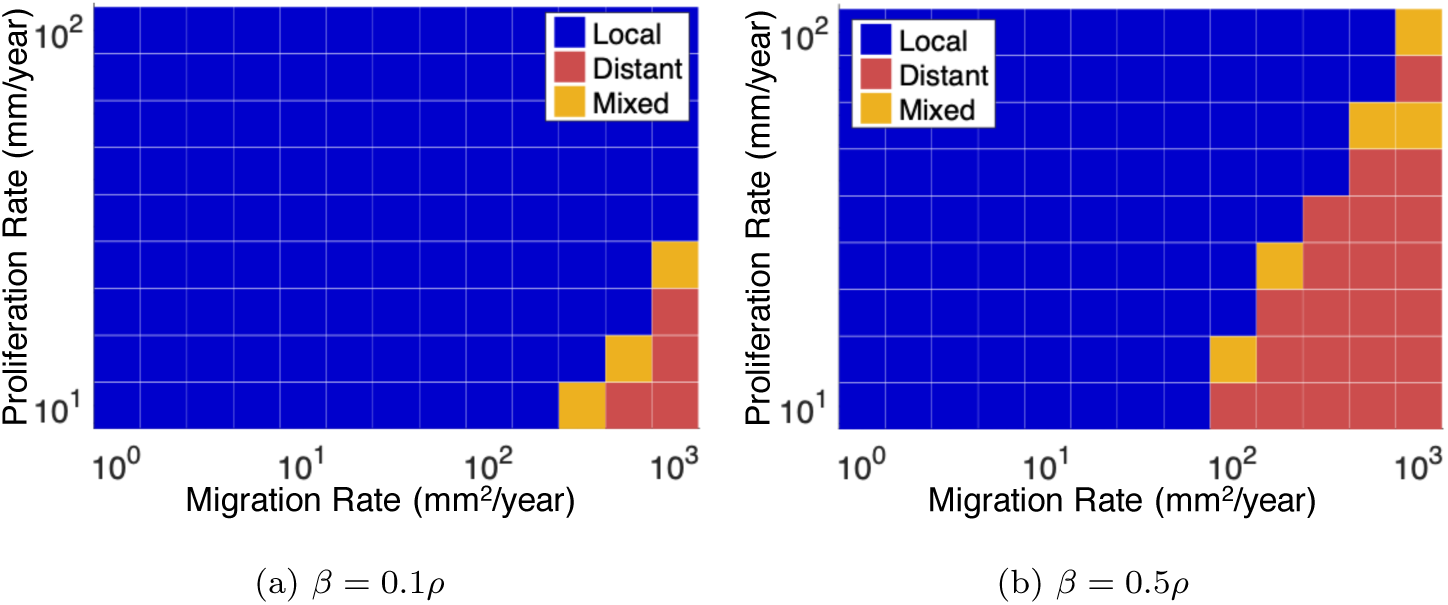
Recurrence location classified for various *D*_*c*_, *ρ, β* and levels of ischemia for *D*_*h*_ = 10*D*_*c*_ for *γ* = 0.05/day. In these simulations, perioperative ischemia was set at 10% of the pre-resection value. We see a larger proportion of local recurrence in these figures compared with those in the main text (see Figure 4).

**Fig. 13:**
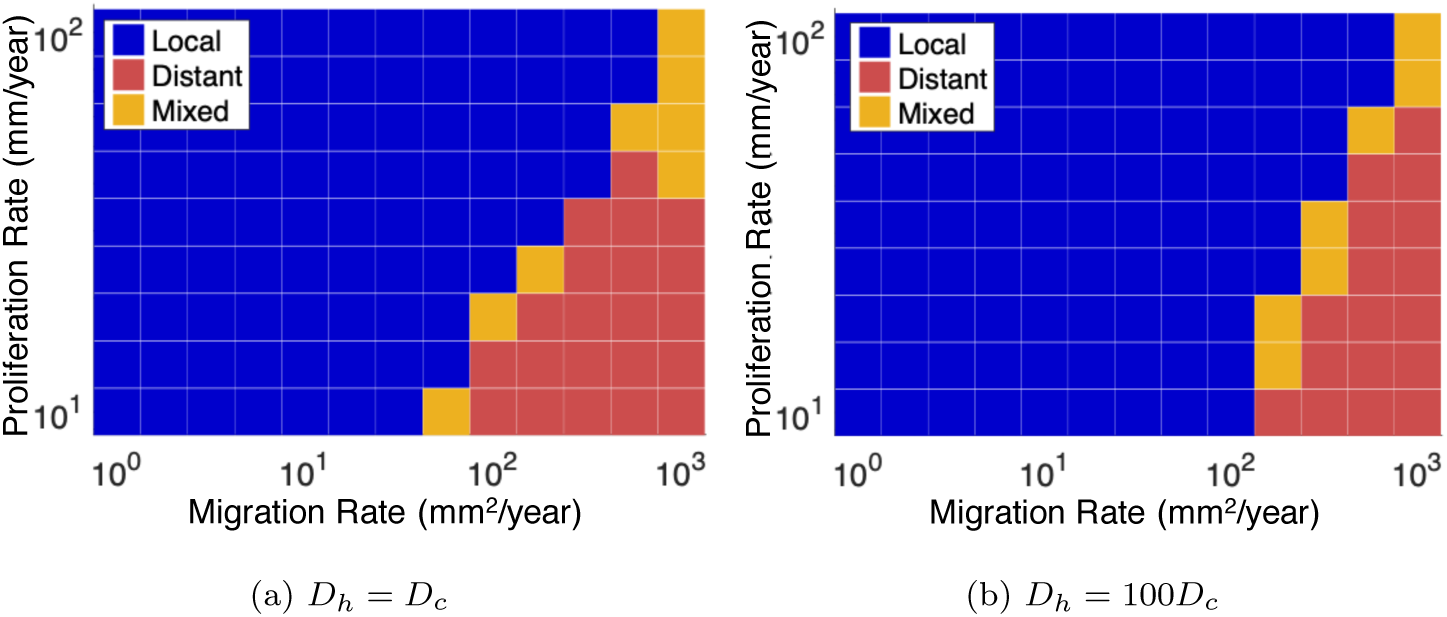
Recurrence location classified for various *D*_*c*_, *ρ, β* and levels of ischemia for *D*_*h*_ = 10*D*_*c*_ for *γ* = 0.005/day. In these simulations, perioperative ischemia was set at 10% of the pre-resection value. We see a larger proportion of local recurrence in these figures compared with those in the main text (see Figure 5).

**Fig. 14:**
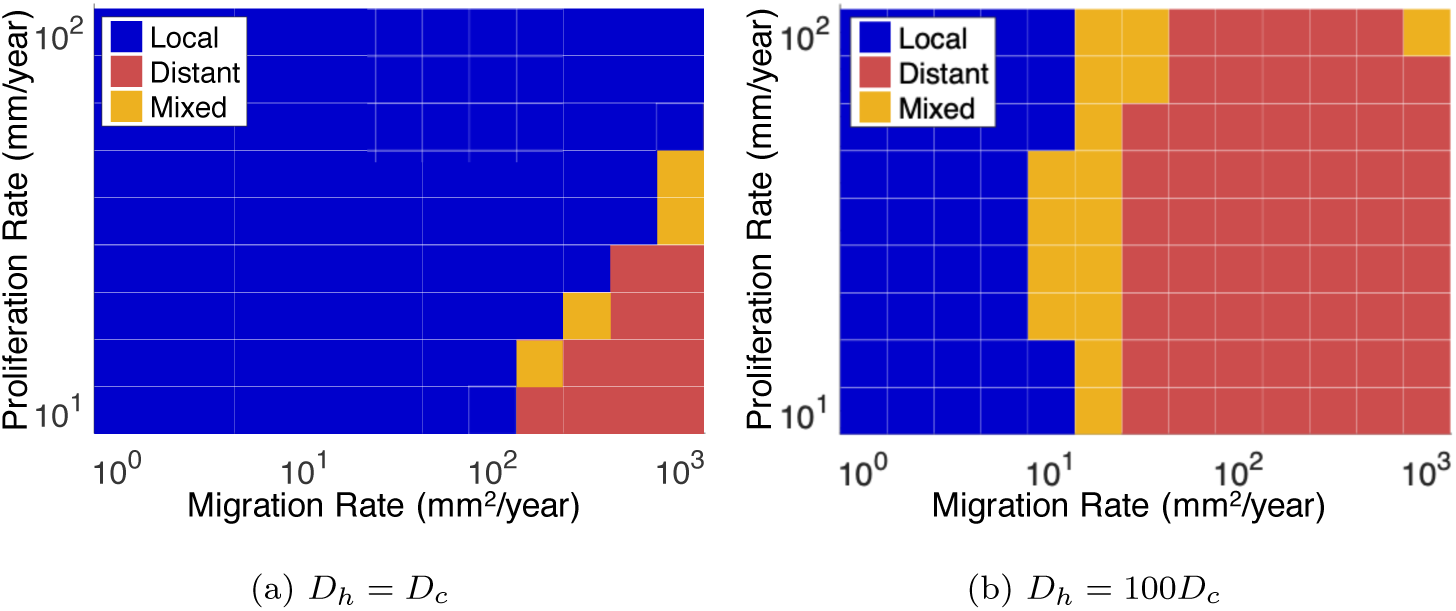
Recurrence location classified for various *D*_*c*_, *ρ* and *D*_*h*_ levels for *β* = 0.5*ρ* and *γ* = 0.05/day. In these simulations, perioperative ischemia was set at 10% of the pre-resection value. We see a larger proportion of local recurrence in these figures compared with those in the main text (see Figure 6).

## 4. Discussion

Through mathematical modeling, we have found a possible mechanism for distal GBM recurrence in response to ischemia. If the tumor has an invasive phenotype, it can remain unimageable on simulated T1Gd MRI as it travels through the ischemic region (using our assumed threshold of 80% total cell density). Once it reaches healthy intact vasculature, it will return to a normoxic phenotype and proliferate to an imageable density outside of the ischemic region before it does so next to the cavity wall. We see that the switching rate from normoxic cells to hypoxic cells plays a role in this behavior, increasing this rate leads to more distantly recurring tumors within the parameter range of *D*_*c*_ and *ρ* that we have used (Figure 4). Conversely, increasing the recovery rate from a hypoxic cell phenotype to a normoxic cell phenotype leads to less distantly recurring tumors (Figure 5). We also note that an increase in the rate of hypoxic cell migration relative to normoxic cell migration promotes distantly recurring tumors (Figure 6).

The migratory nature of GBM cells is a key limitation of conventional treatment efficacy and contributes to tumor recurrence [34]. The dependency of distant recurrence on cell migration shown in these simulations suggests that the use of anti-migratory drugs may reduce the cases of distal recurrence, especially in instances of perioperative ischemia. However, this result is purely theoretical at this stage. These results may also be suggestive of tumor response to hypoxic conditions more generally. Future work may explore patient data to compare pre-operative infiltration patterns with distance to recurrence.

We have shown the intensity of the ischemia plays a role in the observed simulated recurrence patterns (see Appendix Figures 12 - 14). A reduction in functional vasculature to 10% of its pre-resection value does not promote distant recurrence for as many values of *D*_*c*_ and *ρ* as the lower value of 1%. As the value of 10% does not promote quite as much hypoxia in the ischemic region, more simulations are able to reach a T1Gd imageable density inside this region and recur locally.

Furthermore, we have shown that similar simulated recurrence patterns occur in a second tumor location, that was located closer to the surface of the brain than the first and in a more functionally-important region. We also simulated the resection of this second location at a smaller size, yet saw similar patterns of recurrence as the first location. This suggests that the observed model behavior is not simply a function of tumor location or the geometries that we chose for the first resection and ischemic regions. We also ran this simulation in a coronal view of the brain to highlight that this recurrence behavior can occur in any plane. In the future, we can bypass this by moving the model to a more realistic 3D space. Simulated T1Gd MRI volumes are inhibited within the ischemic region, which may explain why diffuse as well as distant recurrences are observed in patients with perioperative ischemia. If the hypoxic cell phenotype were maintained following exposure to ischemia, the tumor as a whole could remain more diffuse in a clinical sense of a large T2 volume relative to T1Gd. Utilizing the PIHNA model may be a useful tool in our effort to understand patterns of recurrence in GBM and understanding the role of ischemia in recurrence and growth patterns more broadly.

## Acknowledgements

The authors gratefully acknowledge funding from the National Cancer Institute (R01CA164371, U54CA193489) and the School of Mathematical Sciences at the University of Nottingham.

## A Recurrence Results of Other PIHNA Simulations

We present the results of PIHNA simulations that were not shown in the main text. The trends in distant recurrence patterns that we observe in the main text all hold in these simulations, supporting our observations regarding *D*_*h*_*/D*_*c*_, *β, γ, D*_*c*_ and *ρ*.

## Footnotes

1 We denote this as *f* to represent fuel for the cells, to avoid reusing *n* which is already assigned to necrotic cells.

2 It is well known that nutrient concentrations in blood (such as glucose concentration) fluctuates throughout a single day, however we are interested in modeling tumor growth over many days and months, so only consider the average nutrient concentration across these daily fluctuations.

## Notes

### Competing Interest Statement

The authors have declared no competing interest.

## References

1. Adeberg, S., König, L., Bostel, T., Harrabi, S., Welzel, T., Debus, J., and Combs, S. E. Glioblastoma recurrence patterns after radiation therapy with regard to the sub-ventricular zone. International Journal of Radiation Oncology* Biology* Physics 90, 4 (2014), 886–893.

2. Ansarizadeh, F., Singh, M., and Richards, D. Modelling of tumor cells regression in response to chemotherapeutic treatment. Applied Mathematical Modelling 48 (2017), 96–112.

3. Armento, A., Ehlers, J., Schötterl, S., and Naumann, U. Molecular mechanisms of glioma cell motility. Exon Publications (2017), 73–93.

4. Barazzuol, L., Burnet, N. G., Jena, R., Jones, B., Jefferies, S. J., and Kirkby, N. F. A mathematical model of brain tumour response to radiotherapy and chemotherapy considering radiobiological aspects. Journal of theoretical biology 262, 3 (2010), 553–565.

5. Bette, S., Barz, M., Huber, T., Straube, C., Schmidt-Graf, F., Combs, S. E., Delbridge, C., Gerhardt, J., Zimmer, C., Meyer, B., et al. Retrospective analysis of radiological recurrence patterns in glioblastoma, their prognostic value and association to postoperative infarct volume. Scientific reports 8, 1 (2018), 4561.

6. Bette, S., Wiestler, B., Kaesmacher, J., Huber, T., Gerhardt, J., Barz, M., Delbridge, C., Ryang, Y.-M., Ringel, F., Zimmer, C., et al. Infarct volume after glioblastoma surgery as an independent prognostic factor. Oncotarget 7, 38 (2016), 61945.

7. Boujelben, A., Watson, M., McDougall, S., Yen, Y.-F., Gerstner, E. R., Catana, C., Deisboeck, T., Batchelor, T. T., Boas, D., Rosen, B., et al. Multimodality imaging and mathematical modelling of drug delivery to glioblastomas. Interface Focus 6, 5 (2016), 20160039.

8. Burnet, N., Jena, R., Jefferies, S., Stenning, S., and Kirkby, N. Mathematical modelling of survival of glioblastoma patients suggests a role for radiotherapy dose escalation and predicts poorer outcome after delay to start treatment. Clinical Oncology 18, 2 (2006), 93–103.

9. Chamberlain, M. Radiographic patterns of relapse in glioblastoma. Journal of Neurooncology 101, 2 (2011), 319–323.

10. Cocosco, C. A., Kollokian, V., Kwan, R. K.-S., Pike, G. B., and Evans, A. C. Brainweb: Online interface to a 3d mri simulated brain database. In NeuroImage (1997), Citeseer.

11. Collins, D. L., Zijdenbos, A. P., Kollokian, V., Sled, J. G., Kabani, N. J., Holmes, C. J., and Evans, A. C. Design and construction of a realistic digital brain phantom. IEEE transactions on medical imaging 17, 3 (1998), 463–468.

12. Curtin, L., Hawkins-Daarud, A., Van Der Zee, K. G., Swanson, K. R., and Owen, M. R. Speed switch in glioblastoma growth rate due to enhanced hypoxia-induced migration. Bulletin of Mathematical Biology 82, 3 (2020), 1–17.

13. de Rioja, V. L., Isern, N., and Fort, J. A mathematical approach to virus therapy of glioblastomas. Biology direct 11, 1 (2016), 1.

14. Delbeke, D., Meyerowitz, C., Lapidus, R., Maciunas, R., Jennings, M., Moots, P., and Kessler, R. Optimal cutoff levels of f-18 fluorodeoxyglucose uptake in the differentiation of low-grade from high-grade brain tumors with pet. Radiology 195, 1 (1995), 47–52.

15. Frieboes, H. B., Lowengrub, J. S., Wise, S., Zheng, X., Macklin, P., Bearer, E. L., and Cristini, V. Computer simulation of glioma growth and morphology. Neuroimage 37 (2007), S59–S70.

16. Gallaher, J. A., Massey, S. C., Hawkins-Daarud, A., Noticewala, S. S., Rockne, R. C., Johnston, S. K., Gonzalez-Cuyar, L., Juliano, J., Gil, O., Swanson, K. R., et al. From cells to tissue: How cell scale heterogeneity impacts glioblastoma growth and treatment response. PLoS computational biology 16, 2 (2020), e1007672.

17. Harpold, H., Alvord, E., and Swanson, K. The evolution of mathematical modeling of glioma proliferation and invasion. Journal of Neuropathology & Experimental Neurology 66, 1 (2007), 1–9.

18. Hawkins-Daarud, A., Rockne, R., Anderson, A., and Swanson, K. Modeling tumor-associated edema in gliomas during anti-angiogenic therapy and its impact on imageable tumor. Frontiers in Oncology 3 (2013), 66.

19. Keunen, O. and Johansson, M., Oudin, A., Sanzey, M., Rahim, S., Fack, F., Thorsen, F., Taxt, T., Bartos, M., Jirik, R., et al. Anti-vegf treatment reduces blood supply and increases tumor cell invasion in glioblastoma. Proceedings of the National Academy of Sciences 108, 9 (2011), 3749–3754.

20. Kwan, R. K.-S., Evans, A. C., and Pike, G. B. An extensible mri simulator for post-processing evaluation. In Visualization in biomedical computing (1996), Springer, pp. 135–140.

21. Kwan, R.-S., Evans, A. C., and Pike, G. B. Mri simulation-based evaluation of image-processing and classification methods. IEEE transactions on medical imaging 18, 11 (1999), 1085–1097.

22. Leder, K., Pitter, K., LaPlant, Q., Hambardzumyan, D., Ross, B. D., Chan, T. A., Holland, E. C., and Michor, F. Mathematical modeling of pdgf-driven glioblastoma reveals optimized radiation dosing schedules. Cell 156, 3 (2014), 603–616.

23. Liberti, M. V., and Locasale, J. W. The warburg effect: how does it benefit cancer cells? Trends in biochemical sciences 41, 3 (2016), 211–218.

24. Louis, D., Ohgaki, H., Wiestler, O., and Cavenee, W. WHO Classification of Tumours of the Central Nervous System, Revised. Fourth Edition. International Agency for Research on Cancer, 2016.

25. Macklin, P., and Lowengrub, J. Nonlinear simulation of the effect of microenvironment on tumor growth. Journal of theoretical biology 245, 4 (2007), 677–704.

26. Martínez-González, A., Calvo, G., Romasanta, L., and Pérez-García, V. Hypoxic cell waves around necrotic cores in glioblastoma: a biomathematical model and its therapeutic implications. Bulletin of Mathematical Biology 74, 12 (2012), 2875–2896.

27. Neufeld, Z., von Witt, W., Lakatos, D., Wang, J., Hegedus, B., and Czirok, A. The role of allee effect in modelling post resection recurrence of glioblastoma. PLoS computational biology 13, 11 (2017), e1005818.

28. Pardo, R., Martinez-Gonzalez, A., and Perez-Garcia, V. M. Nonlinear ghost waves accelerate the progression of high-grade brain tumors. Communications in Nonlinear Science and Numerical Simulation 39 (2016), 360–380.

29. Raza, S. M., Lang, F. F., Aggarwal, B. B., Fuller, G. N., Wildrick, D. M., and Sawaya, R. Necrosis and glioblastoma: a friend or a foe? a review and a hypothesis. Neurosurgery 51, 1 (2002), 2–13.

30. Rockne, R. C., Trister, A. D., Jacobs, J., Hawkins-Daarud, A. J., Neal, M. L., Hendrickson, K., Mrugala, M. M., Rockhill, J. K., Kinahan, P., Krohn, K. A., et al. A patient-specific computational model of hypoxia-modulated radiation resistance in glioblastoma using 18f-fmiso-pet. Journal of the Royal Society Interface 12, 103 (2015), 20141174.

31. Roniotis, A., Sakkalis, V., Tzamali, E., Tzedakis, G., Zervakis, M., and Marias, K. Solving the pihna model while accounting for radiotherapy. In Advanced Research Workshop on In Silico Oncology and Cancer Investigation-The TUMOR Project Workshop (IARWISOCI), 2012 5th International (2012), IEEE, pp. 1–4.

32. Rutter, E. M., Stepien, T. L., Anderies, B. J., Plasencia, J. D., Woolf, E. C., Scheck, A. C., Turner, G. H., Liu, Q., Frakes, D., Kodibagkar, V., et al. Mathematical analysis of glioma growth in a murine model. Scientific reports 7, 1 (2017), 1–16.

33. Scribner, E., Saut, O., Province, P., Bag, A., Colin, T., and Fathallah-Shaykh, H. M. Effects of anti-angiogenesis on glioblastoma growth and migration: model to clinical predictions. PLoS One 9, 12 (2014), e115018.

34. Silbergeld, D., and Chicoine, M. Isolation and characterization of human malignant glioma cells from histologically normal brain. Journal of Neurosurgery 86, 3 (1997), 525–531.

35. Stark, A. M., van de Bergh, J., Hedderich, J., Mehdorn, H. M., and Nabavi, A. Glioblastoma: clinical characteristics, prognostic factors and survival in 492 patients. Clinical neurology and neurosurgery 114, 7 (2012), 840–845.

36. Stein, A. M., Demuth, T., Mobley, D., Berens, M., and Sander, L. M. A mathematical model of glioblastoma tumor spheroid invasion in a three-dimensional in vitro experiment. Biophysical journal 92, 1 (2007), 356–365.

37. Stupp, R., Hegi, M., Mason, W., van den Bent, M., Taphoorn, M., Janzer, R., Ludwin, S., Allgeier, A., Fisher, B., Belanger, K., et al. Effects of radiotherapy with concomitant and adjuvant temozolomide versus radiotherapy alone on survival in glioblastoma in a randomised phase iii study: 5-year analysis of the eortc-ncic trial. The Lancet Oncology 10, 5 (2009), 459–466.

38. Subramanian, S., Gholami, A., and Biros, G. Simulation of glioblastoma growth using a 3d multispecies tumor model with mass effect. Journal of mathematical biology 79, 3 (2019), 941–967.

39. Swan, A., Hillen, T., Bowman, J., and Murtha, A. A patient-specific anisotropic diffusion model for brain tumour spread. Bulletin of Mathematical Biology 80, 5 (2018), 1259–1291.

40. Swanson, K. Mathematical Modeling of the Growth and Control of Tumors. PhD thesis, University of Washington, 1999.

41. Swanson, K., Alvord, Jr, E., and Murray, J. A quantitative model for differential motility of gliomas in grey and white matter. Cell Prolif 33, 5 (Oct 2000), 317–29.

42. Swansonxs, K., Bridge, C., Murray, J., and Alvord, E. Virtual and real brain tumors: using mathematical modeling to quantify glioma growth and invasion. Journal of the Neurological Sciences 216, 1 (2003), 1–10.

43. Swanson, K., Rockne, R., Claridge, J., Chaplain, M., Alvord, E., and Anderson, A. Quantifying the role of angiogenesis in malignant progression of gliomas: in silico modeling integrates imaging and histology. Cancer Research 71, 24 (2011), 7366–7375.

44. Swanson, K., Rostomily, R., and Alvord, E. A mathematical modelling tool for predicting survival of individual patients following resection of glioblastoma: a proof of principle. British Journal of Cancer 98, 1 (2008), 113–119.

45. Thiepold, A., Luger, S., Wagner, M., Filmann, N., Ronellenfitsch, M., Harter, P., Braczynski, A., Dützmann, S., Hattingen, E., Steinbach, J., et al. Perioperative cerebral ischemia promote infiltrative recurrence in glioblastoma. Oncotarget 6, 16 (2015), 14537.

46. Warburg, O. The metabolism of carcinoma cells. The Journal of Cancer Research 9, 1 (1925), 148–163.

47. Yamaguchi, T., Kanno, I., Uemura, K., Shishido, F., Inugami, A., Ogawa, T., Murakami, M., and Suzuki, K. Reduction in regional cerebral metabolic rate of oxygen during human aging. Stroke 17, 6 (1986), 1220–1228.

48. Yang, Y., Hou, L., Li, Y., Ni, J., and Liu, L. Neuronal necrosis and spreading death in a drosophila genetic model. Cell death & disease 4, 7 (2013), e723.

49. Zagzag, D., Lukyanov, Y., Lan, L., Ali, M. A., Esencay, M., Mendez, O., Yee, H., Voura, E. B., and Newcomb, E. W. Hypoxia-inducible factor 1 and vegf upregulate cxcr4 in glioblastoma: implications for angiogenesis and glioma cell invasion. Laboratory Investigation 86, 12 (2006), 1221.

50. Zuniga, R., Torcuator, R., Jain, R., Anderson, J., Doyle, T., Ellika, S., Schultz, L., and Mikkelsen, T. Efficacy, safety and patterns of response and recurrence in patients with recurrent high-grade gliomas treated with bevacizumab plus irinotecan. Journal of Neuro-oncology 91, 3 (2009), 329.

